# STRAIGHTMORPH: A Voice Morphing Tool for Research in Voice Communication Sciences

**DOI:** 10.1101/2024.06.04.597359

**Authors:** P Belin, H Kawahara

## Abstract

The purpose of this paper is to make easily available to the scientific community an efficient voice morphing tool called STRAIGHTMORPH and provide a short tutorial on its use with examples. STRAIGHTMORPH consists of a set of Matlab functions allowing to generate high-quality, parametrically-controlled morphs of an arbitrary number of voice samples. A first step consists in extracting an ‘mObject’ for each voice sample, with accurate tracking of the fundamental frequency contour and manual definition of Time and Frequency anchors corresponding across samples to be morphed. The second step consists in parametrically combining the mObjects to generate novel synthetic stimuli, such as gender, identity or emotion continua, or random combinations. Although STRAIGHTMORPH has been designed for human voices it can work well with other types of sounds such as non-human primate vocalizations.

## Why Voice Morphing?

The ability to parametrically manipulate the physical characteristics of basic sensory stimuli – e.g., the direction of a moving bar on a screen, or the frequency of a pure tone—has played a major role in experimental psychology. But what is easy to perform for simple synthetic stimuli becomes much harder with complex naturalistic stimuli such as faces or voices.

The availability of face morphing software has resulted in important advances in our understanding of the perceptual and cerebral representations of face identity and emotion. Thanks to these tools, researchers can generate synthetic stimuli that preserve essential perceptual aspects of faces and that can be parametrically controlled. For instance, one can generate a continuum of faces between 2 identities, or between one identity and an average face, such that the physical distance between any two consecutive faces is exactly similar at any point of the continuum, allowing a dissociation of physical vs. perceptual differences between stimuli (Koyano et al., 2021; D. A. Leopold, Bondar, & Giese, 2006; D.A. Leopold, O’Toole, Vetter, & Blanz, 2001; Morris et al., 1996; Rotshtein, Henson, Treves, Driver, & Dolan, 2005).

## Legacy STRAIGHT

A similar implement thankfully exists in the voice modality: the open-source Matlab package Legacy STRAIGHT has been developed by the second author as a speech vocoder allowing high-quality transformations of the speech signal (Kawahara & Matsui, 2003; Kawahara & Morise, 2024) and its use by an growing number of researchers yields exciting results (Bruckert et al., 2010; Charest, Pernet, Latinus, Crabbe, & Belin, 2013; Latinus, McAleer, Bestelmeyer, & Belin, 2013; Nussbaum, von Eiff, Skuk, & Schweinberger, 2022; Skuk et al., 2020 ; von Eiff, Fruhholz, Korth, Guntinas-Lichius, & Schweinberger, 2022) }. The vocoder tools decompose input speech signals into sets of vocoder parameters. They are a) the spectral envelope (filter parameter), b) the fundamental frequency (f0), and c) the aperiodicity (time-frequency map of ratio between random and periodic components). To achieve this, STRAIGHT performs an instantaneous pitch-adaptive spectral smoothing based on robust f0 extraction for pitch-synchronous estimation of the spectral profile. STRAIGHT generates a ‘smoothed ‘ spectrogram using a complementary set of windows designed by the extracted f0 and inverse filtering in spline space. The extracted f0 is also applied to separate the periodic vs. aperiodic parts of the vocal signal. The periodic and aperiodic parts of the signal can then be recombined and resynthesized, generally yielding a high-quality synthetic version of the original stimulus. They can also be independently and parametrically manipulated or combined across several samples before resynthesis, yielding novel synthetic stimuli. Although it has been developed primarily as a high-quality speech vocoder, STRAIGHT also works very well for manipulation of other types of vocal information such as identity or emotion, and can work well with vocalizations from other species (Chakladar, Logothetis, & Petkov, 2008; Furuyama, Kobayasi, & Riquimaroux, 2017)

The purpose of the present paper is to make easily available to the scientific community an efficient voice morphing tool called STRAIGHTMORPH and provide a short tutorial on its use with examples. STRAIGHTMORPH consists of Matlab functions from the open source package Legacy STRAIGHT, plus functions developed in the first author’s Voice Neurocognition Laboratory particularly

### ExtractMObject and VoiceMultiMorph

## STRAIGHTMORPH

STRAIGHTMORPH requires a recent version of Matlab with the Signal Processing toolbox. All files can be accessed for download at https://github.com/pascalbelin/STRAIGHTMORPH. The folder ‘scripts’ should be added to the Matlab path. The TUTORIAL folder contains example sounds and mObjects to be used with the script STRAIGHTMORPH_tutorial.m

### Step 1 – Extracting mObjects

The function **ExtractMObject.m** calls STRAIGHT scripts to analyze a sound and generate an ‘mObject’ for this sound. MObjects have been devised by the second author as a convenient way to store all the information concerning each sound to be analyzed, resynthesized and combined with other sounds. They consist of a Matlab structure with 16 different fields. The most important fields are:

i. mObject.**waveform** containing the stimulus waveform;
ii. mObject.**F0** containing a vector of length N equal to the duration of the stimulus in milliseconds, with f0 estimates every millisecond;
iii. mObject.**spectrogram**, containing a 513 (frequencies) x N (millisecond time points) matrix containing the smoothed spectrogram of the stimulus’s periodic component;
iv. mObject.**aperiodicityIndex** containing a matrix of similar size as the spectrogram but containing an time-frequency map of ratio between random and periodic components;
v. mObject.**anchorTimeLocation** and mObject.**anchorFrequency** containing information on the time and frequencies, respectively, of the Time and Frequency anchors used to combine several mObjects.

ExtractMObject invites **manual user intervention** at two critical steps (also detailed in the script):

i. **F0 estimation**. Estimation of the time varying fundamental frequency (f0) is a critical step in extraction of the mObject and largely determines the quality of resynthesis and then morphing. The robust f0 extractor in STRAIGHT works well in most cases, but in some cases of e.g. noisy or highly aperiodic sounds the automatic procedure yields unsatisfying f0 trajectories, with e.g. interruptions or shifts to higher harmonics in some portions of the signal. ExtractMObject offers the option to manually limit the frequency range in which f0 candidates are analyzed to try and correct unwanted discontinuities (Fig. 1B,C). Upon calling the function a new window opens displaying a time-frequency graph with different f0 candidates. <u>Note that the mouse should stay on that window to avoid problems when clicking</u>. The f0 trajectory selected by the automatic procedure will be highlighted in green. If the result of this automatic f0 detection looks correct, without unnecessary interruptions, the user can skip the manual adjustment step by pressing the Esc key and move to the next step. If, however, the result of the automatic f0 detection looks unsatisfying, the user clicks on the figure to manually define lower and upper frequency constraints. First the lower boundary is defined by clicking with the left button to add landmarks and with the right button to finish boundary definition. Then the same has to be done for the upper frequency boundary. The F0 detection process will be restarted by STRAIGHT taking into account the manually defined constraints.
ii. **Definition of Time and Frequency anchors**. A crucial step in face morphing consists in defining landmarks to be put in correspondence across faces, i.e., eyes with eyes and mouth with mouth. Similarly, STRAIGHT makes use of spectro-temporal anchors to constrain the interpolation between different stimuli. Defining accurately time and frequency anchors is of crucial importance for high-quality morphing. Time anchors usually indicate beginning and end of the sound, phoneme temporal boundaries and important formant transitions. Frequency anchors usually indicate formant central frequencies. <u>The number of time axes should be the same for all sounds to be morphed together</u>. The number of frequency anchors can be different for different time axes; yet <u>for each time axis, the number of frequency anchors has to be consistent across all the sounds</u>.

**Figure 1.**
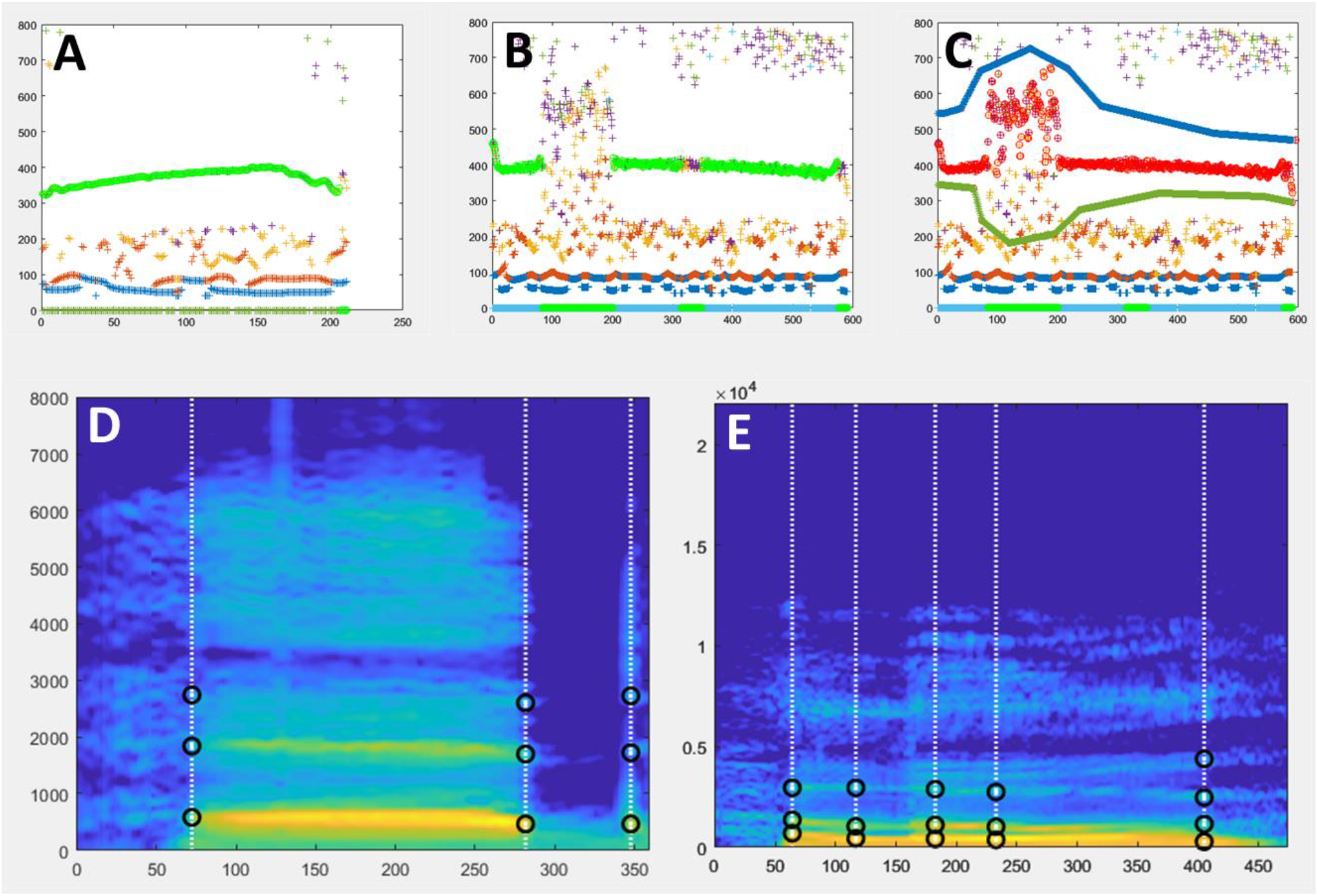
mObject extraction. **A**. Example of successful automatic f0 extraction using ExtractMObject. The diagram (X axis: time (ms); Y axis: frequency(Hz)) shows identified f0 candidates as coloured circles. F0 candidates selected by the automatic procedure are highlighted in green. **B**. Example of problematic automatic f0 extraction. The green f0 contour is interrupted at several instances which will cause artefacts at resynthesis. This problem can be minimized using manual edition of f0 search range. **C**. Example in B for which f0 search range has been restricted manually to minimize interruptions. The new selected f0 contour is shown in red. **D, E**. Examples of Time and Frequency anchors (black circles) shown on the smoothed spectrograms of two mObjects to be used in examples below.

ExtractMObject provides a graphical interface to manually set Time and Frequency anchors (these values can also be edited in the anchorTimeLocation and anchorFrequency fields of the mObject structure). Directly after the f0 extraction stage, a smoothed spectrogram of the analyzed stimulus is displayed full screen. The user has to first define a Time location by clicking with the right mouse button at the appropriate temporal location below the x (time) axis. This will display a vertical time line on which candidate Frequency anchors (local energy maxima) appear at different frequencies as black circles. The user then defines one or more frequency anchors on this time axis by clicking at the chosen frequency with the left button. This process can be repeated as necessary to create new Time locations and selecting new Frequency anchors. The user can press Esc to start the definition process again or Enter to finalize MObject extraction.

Figure 1D shows the smoothed spectrogram of an mObject with its Time and Frequency anchors shown as black circles: the word ‘head’ spoken by a male speaker (M048eh). Time anchors are positioned at onset and offset of the voiced part of the first syllable, and at plosive onset; Frequency anchors are positioned on the first 3 formants. Anchors can be more complex such those shown for the word ‘Hello’ spoken a female speaker (Fig 1E).

Once an mObject has been generated, it can be first be inspected using the STRAIGHT function displayMObject (Code in Appendix lines xx). Then <u>it is crucial to check its quality by resynthesizing it</u> and listening to the resynthesized version compared to the original. When MObject extraction works well, the difference between the original and resynthesized versions should be barely perceptible, if at all. A noticeable difference between the two, likely arising from a problem with the f0 extraction step, indicates poor MObject extraction likely to contaminate all morphed stimuli and that should preferably be redone.

### Step 2 – Combining mObjects

The function **VoiceMultiMorph** pilots the morphing of an arbitrary number of MObjects. Its general functioning is explained below then illustrated by different examples. All MObjects to be morphed are processed in a symmetrical way, without intermediate morphing stages. The mObjects to be morphed are first loaded in a cell array of structure. A critical step then consists of defining the *morphing rates* (‘mRates’) for each sound. <u>mRates can be thought of as the coordinates of the sound to be generated in a multidimensional voice space with as many dimensions as original mObjects</u>. They consist of a structure with 5 fields, each of them being a vector with as many elements as MObjects indicating the weight of each MObject for interpolation of that particular field. The 5 fields are: (i) F0: mRates vector for f0; (ii) spectralamplitude : mRates vector for spectral amplitude; (iii) aperiodicity: mRates vector for aperiodicity index; (iv) time: mRates vector for Time anchors; (v) frequency: mRates vector for Frequency anchors. <u>For each of the 5 fields the sum of all mRates must equal one for that field</u>. mRates for a given sound can be different across the 5 fields (cf. ‘Parameter-specific morphing’ below).

Default settings for interpolation based on mRates are logarithmic for f0, spectralamplitude and Frequency anchors; linear for Time anchors and aperiodicity indices (which are on a dB scale). Calling the function Voicemultimorph will generate a new mObject (‘mObjectM’) based on the mRates provided for the different MObjects, which can then be examined and resynthesized as any other mObject.

## MORPHING EXAMPLES

### Morphing 2 Mobjects

The examples below details the interpolation of recordings of a male speaker and a female speaker uttering the same word from an excellent database of American English Vowels (Hillenbrand, Getty, Clark, & Wheeler, 1995) but can be applied to any other pair of mObjects such as recordings of different emotional expressions from the same speaker.

#### Generating the average of two mObjects

To generate an average of the mObject1 (Fig. 2A) and mObject2 (Fig. 2C), a rate of 0.5 is set for each mObject, such that each the 5 mRates fields equals [0.5 0.5]. Running VoiceMultiMorph generates the new mObjectM which can then be synthesized, listened to, saved if necessary and inspected. Figure 2b shows the smoothed spectrogram of the 2-voice average. Note that since the Frequency anchors of the two original mObjects in this example correspond to frequencies of their first 3 formants, the Frequency anchors of the newly generated mObjectM also provide its 3 formant frequencies.

**Figure 2.**
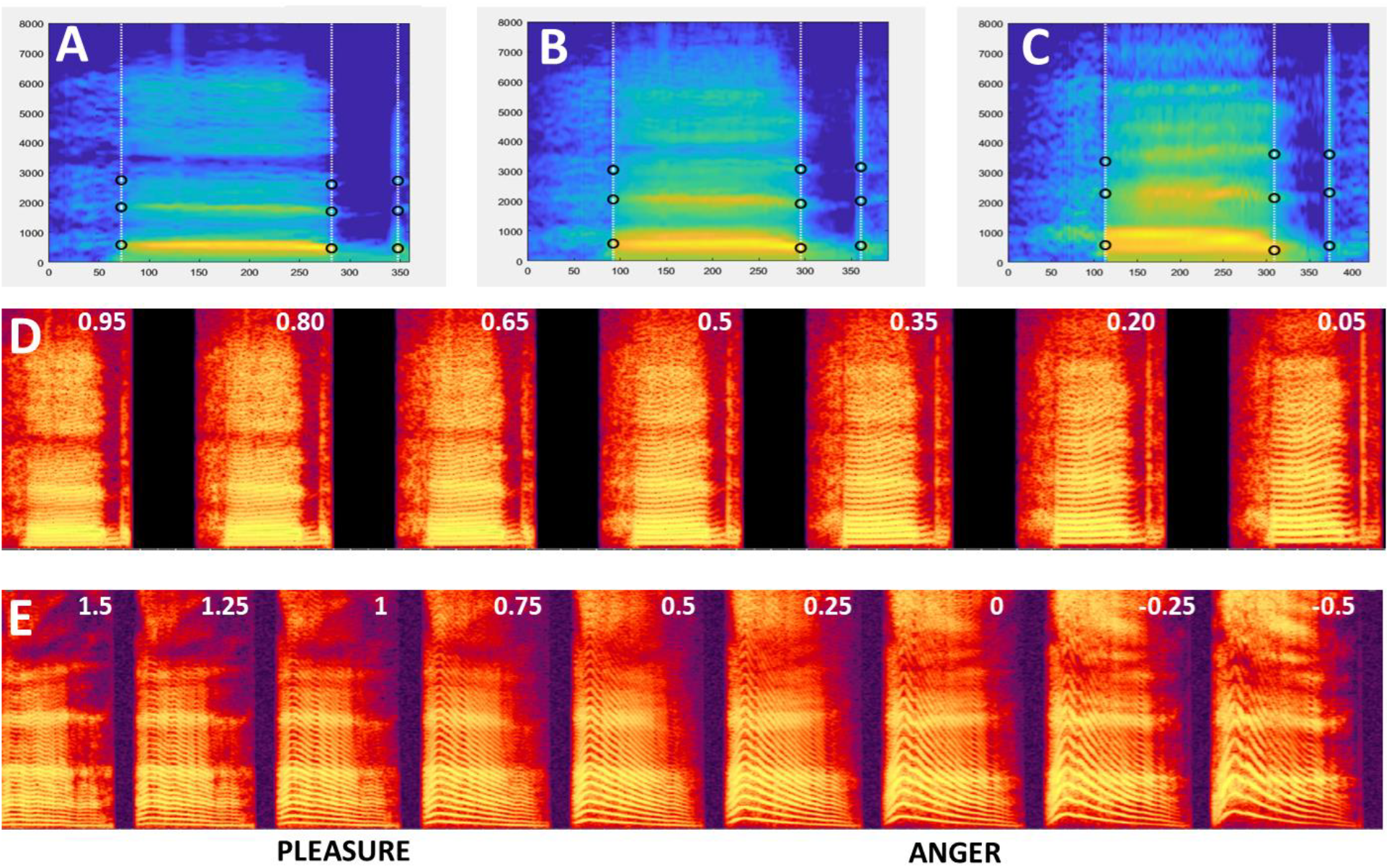
Morphing 2 stimuli. **A-C**. Smoothed spectrograms with Time and Frequency anchors of 3 Mobjects. **A:** M48eh. **C:** W33eh. Note the similar definition of Time and Frequency anchors. B: Average of M48eh and W33eh. **D**. Spectrogram of the continuum generated between M48eh and W33eh using values of k =[0.95:-0.15:0.05] . Values of k are indicated at the top of each spectrogram. The leftmost stimulus is a 95%/5% mix of M48eh and W33eh, decreasing/increasing in steps of 15% until the rightmost stimulus – 5%/95% M48eh and 95% W33eh. The middle stimulus is the 50%/50% average shown in B. Any pair of consecutive stimuli is separated by the same acoustical distance at any point of the continuum: 15% of the M48eh-W33eh distance. **E**. Pleasure-Anger continuum including caricatures generated using values of k=[1.5:-.25:-0.5].

#### Generating a continuum between two sounds

To generate a continuum between the two sounds, variable rates are set using a parameter k: mRates=[k 1-k], with k between 0 and 1. k represents the weight of the first mObject, such that a value of k=1 (mRates=[1 0]) corresponds to the first mObject, a value of k=0 (mRates=[0 1]) corresponds to the second mObject, and values between 0 and 1 to interpolation between the 2 mObjects. Parameter k can be set at regular intervals between 0 and 1 to generate a series of mObjects regularly spaced between the 2 original mObjects. For instance k=[1:-0.1:0] will generate a series of 11 mObjects in 10% steps between the two original mObjects. Or k=[0.95:-0.15:0.05] will generate 7 mObjects in 15 % steps between a 95%/5% and a 5%/95% morphs – such that each sound of the continuum contains some part of the two original sounds (Figure 2D). Such continua are particularly interesting as <u>they provide series of stimuli with equal acoustical distance between any two consecutive stimuli</u> and can thus be used to dissociate perceptual aspects of a particular voice dimension (here, gender) from the unavoidable physical differences between the stimuli. For instance, the same 15% acoustical difference between 2 consecutive gender morphs has very different perceptual effects depending on whether they are towards the extremes of the continuum, or towards the middle of the continuum where it leads to marked changes in perceived gender (Charest et al., 2013; Pernet & Belin, 2012).

#### Caricaturing

In the continuum example of Figure 2D mRates kept values between 0 and 1, which makes intuitive sense but is not mathematically required. Indeed, while mRates values between 0 and 1 correspond to stimuli generated between mObject1 and mObject2 in the line between the two in multidimensional voice space, values below 0 and above 1 correspond to points beyond the two mObjects on that same line; i.e., caricatures (Calder et al., 2000; Whiting, Kotz, Gross, Giordano, & Belin, 2020).

For instance, mRates = [1.5 -0.5] correspond to a point on the line at 50% of the mObject1-mObject2 distance on the side of mObject1 – in the example above, a voice exaggerating gender difference towards the male voice – while mRates=[-0.5 1.5] will produce and exaggeration of the same magnitude but towards the female voice. So by using mRates=[k 1-k] and k extending beyond 0 and 1 the continuum will be extended on both sides with increasingly strong caricatures . Fig 3E shows the spectrogram of an emotion continuum generated using k=[1.5:-0.1:-0.5] between a pleasure (6_pleasure.wav) and an angry (6_anger.wav) expression from an actor of the Montreal Affective Voices (Belin, Fillion-Bilodeau, & Gosselin, 2008). Note that using mRates values too negative or too much above 1 will likely result in unnatural sounding stimuli…

**Figure 3.**
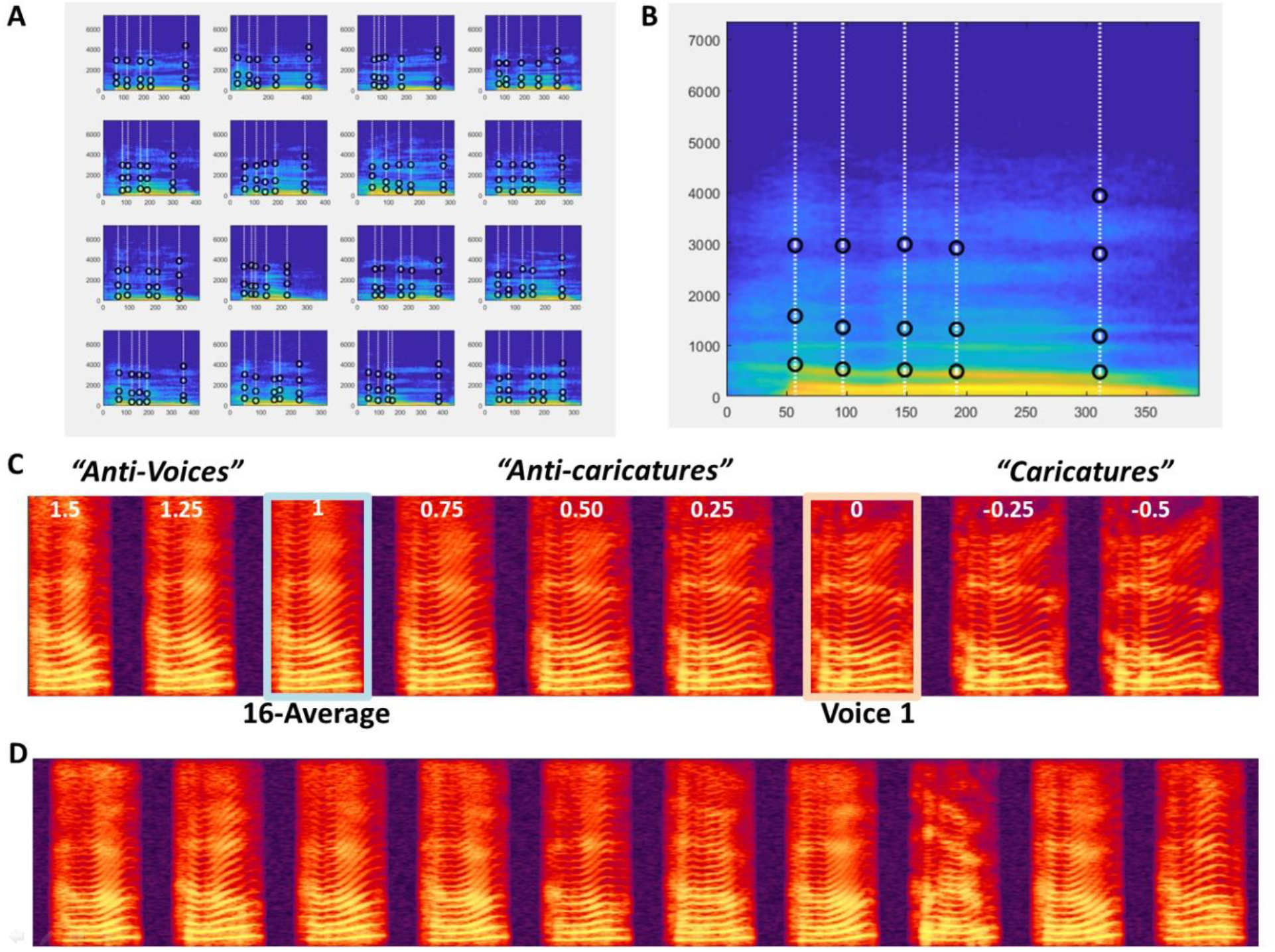
Morphing 16 mObjects. **A**. 16 mObjects of the word Hello spoken by 16 female speakers. Note their similar definition of the Time and Frequency anchors for combined morphing. **B**. mObjectM corresponding to the average of the 16 mObjects. **C**. Spectrogram of Continuum between Voice1 and the 16-Average with negative values of k for Caricatures and values of k greater than 1 for ‘Antivoices’ (Latinus & Belin, 2011). **D**. 10 random ‘Hellos’ generated using random mRates.

#### Parameter-specific morphing

In the examples above all 5 fields of the mRates structure were set to the same values. But it can be desirable in some cases to set some fields to different values than other, what has been called ‘parameter-specific morphing’ (Skuk et al., 2020). For instance, all stimuli of the continuum could be set to the same duration – the average duration of mObjects 1 and 2 – by using mRates.duration=[0.5 0.5] while mRates in the other fields vary with k. Or, the anchorFrequency field could be set to [0.5 0.5] while the others vary with k, to generate a continuum of stimuli all having the formant frequencies of the mObject1-mObject2 average.

### Morphing N MObjects

A significant feature of VoiceMultiMorph is that it treats morphing N>2 mObjects in exactly the same way as for 2 mObjects. Simply, there are N dimensions instead of 2 to the multidimensional voice space defined by the mObjects, and mRates fields consist of a vector of N values (still adding up to 1 for each field) that can be viewed as the coordinates of the new mObjectM in N-dimensional voice space. Rather than averaging in turn all stimuli in successive steps, the user simply defines the desired coordinates in that space, and performs a single generation of mObjectM and resynthesis no matter how many stimuli are combined. In the examples below we consider 16 mObjects generated from recordings of the word ‘Hello’ spoken by 16 female speakers (Fig. 3A) as part of a study on voice personality impressions (McAleer, Todorov, & Belin, 2014). The first step consists in loading the 16 mObjects in a cell array of structures.

#### Generating the N-Average

As for 2 mObjects, to generate the average of N mObjects the mRates vectors are all set to the same value of 1/N, here mRates=*1/16 … 1/16+ or ones(1,16)/16. The mObjectM generated by these mRates values can be viewed as the center of N-dimensional voice space. It has average f0 contour and duration; it has average aperiodicity map, resulting in a very smooth spectrogram (Fig 3B) with high harmonics-to-noise ratio. Such an average stimulus is typically perceived as more attractive than individual voices (Bruckert et al., 2010) and elicits lower activity in the TVAs than individual voices (Bestelmeyer et al., 2011; Latinus et al., 2013)– potentially reflecting a norm-based coding mechanism relative to an internal prototype.

#### Morphing with the N-Average

It can be interesting to interpolate between a single mObject and the N-Average: studies in the face perception domain use this strategy to modulate the ‘identity strength’ of the faces (D.A. Leopold et al., 2001). To do so, one strategy could be to first generate the mObject corresponding to the N-Average, then generate the continuum between the 2 mObjects as in the example above. Another, better strategy, that skips an unnecessary step, is to specify directly the mRates of the desired continuum trajectory in N-dimensional space.

With k=[1.5:-0.1:-0.5] as in the example above, to interpolate between the N-Average and mObject1, using mRates = [k*ones(1,16)/16] + [(1-k)*[1 zeros(1,15)]] or ([ 1-k+k/16 k/16 k/16… k/16+ will generate stimuli on the line in multidimensional space going through mObject1 and the N-Average. K=1 corresponds to the average; k=0 to mObject 1. Values of k between O and 1 correspond to stimuli between the N-Average and mObject1 (‘anti-caricatures’); values of k below 0 correspond to caricatures of mObject1 relative to the average. Values of k above 1 correspond to ‘anti-voices’, i.e. stimuli on the other side of the average relative to mObject1 in voice space, with opposite characteristics, e.g., higher than average f0 if mObject1 has lower f0 than average, etc. (Figure 3C)

Generating continua with the N-average can also be used for instance to take advantage of its high harmonics-to-noise ratio and generate ‘smoother’ or ‘rougher’ versions of mObjects. By using mRates values of *1 0 0 … 0+ for all mRates fields, except for mRates.aperiodicity=*k*ones(1,16)/16+ + [(1-k)*[1 zeros(1,15)]] stimuli will keep all aspects of mObject 1 but for the degree of aperiodic noise, and moving it closer to the average will generate a version of mObject1 with less aperiodicity, which listeners find more attractive (Bruckert et al., 2010)

#### Generating Random Stimuli

The N mObjects can be viewed as the axes of an N-dimensional voice space they define. While all points of that space likely do not correspond to naturalistic voice stimuli – depending of course on the choice of the N mObjects – many points in the region around the average likely do. The N mObjects thus constitute a generative space that can be used to synthesize any point of the space. By applying random weights to each mObject in the mRates (still ensuring the total adds up to 1 per field), it is possible to sample the voice space by generating an arbitrary number of random stimuli. The example in the STRAIGHTMORPH_tutorial.m script applies random weights, with a coefficient that modulates the radius of the hypersphere around the average in which stimuli are generated, in order to generate an arbitrary number of random ‘Hellos’ and keeps the same weights across the 5 fields of the mRates.

### Morphing sounds other than human voice

Finally, STRAIGHTMORPH can be used to morph stimuli other than human voices, provided there is a relatively clear f0. For instance, it is possible to generate a continuum between a human voice and a note played by a musical instrument (Belizaire, Fillion-Bilodeau, Chartrand, Bertrand-Gauvin, & Belin, 2007). Several groups have applied STRAIGHT to manipulation and morphing of vocalization from nonhuman primates (Chakladar et al., 2008; Furuyama et al., 2017). ExtractMObject will work well on these vocalizations provided the periodic content is strong enough but may require adjustment of the default values for stimuli with e.g. high f0. Figure 4A shows an example of identity continuum for macaque ‘Coo’ vocalizations similar to that of Fig 3C. For stimuli with very high f0, such as marmoset vocalizations, good results can be obtained by first transposing stimuli in the speech range by adjusting the sampling rate, then perform morphing, then readjusting the sampling rate of the generated stimuli (Fig. 4B).

**Figure 4.**
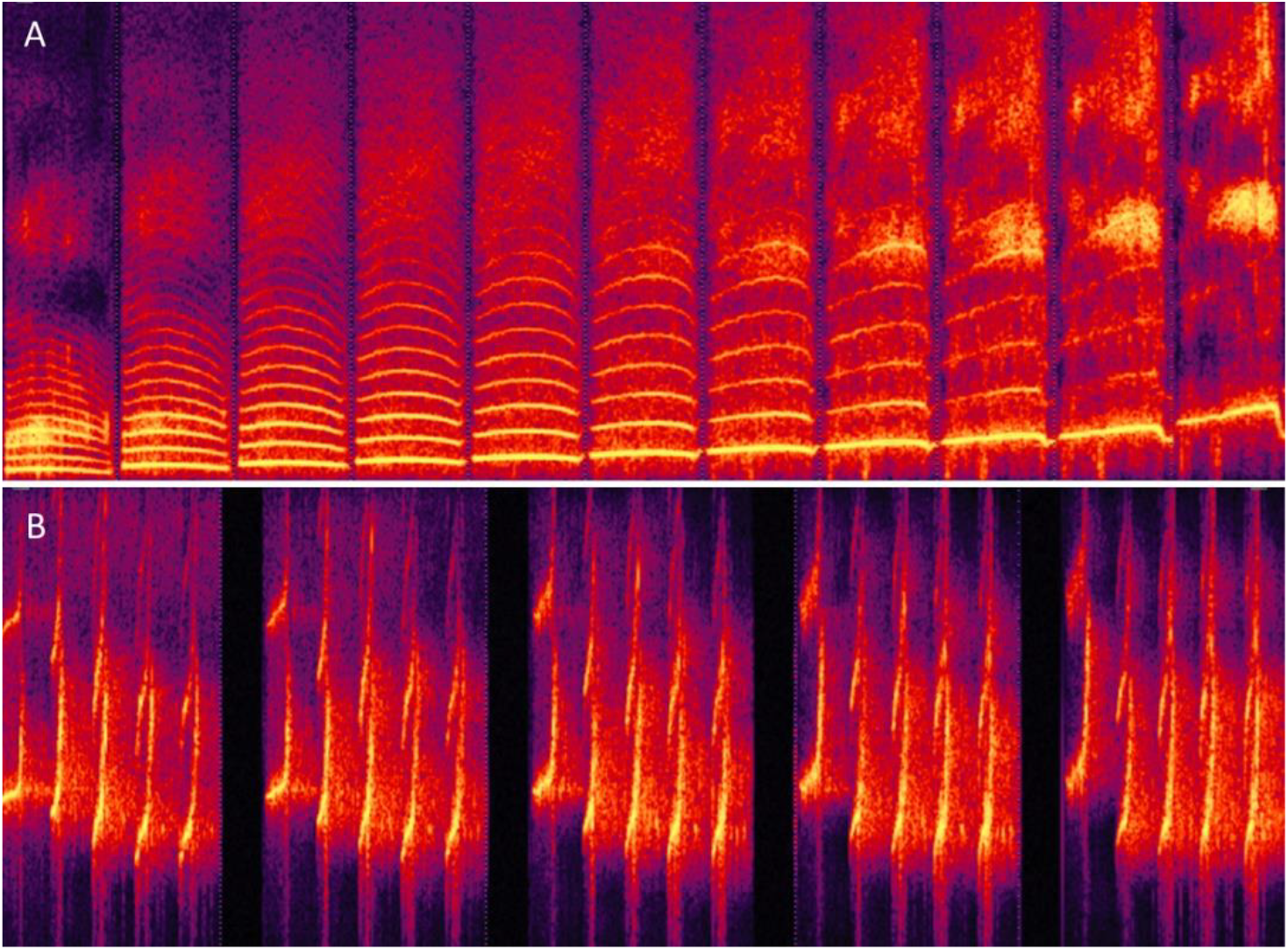
Morphing non-human primate vocalizations. **A**. Continuum between an individual macaque Coo and the average of 16 Coos using values of k similar to Fig. 3C. **B**. Continuum between Twitter calls from 2 individual marmoset monkeys.

## ACKNOWLEDGEMENTS

We thank Dr Julien Rouger and ‘a wee bit’ Dr Marianne Latinus for developing the ExtractMObject and VoiceMultiMorph functions in Glasgow’s Voice Neurocognition Laboratory. We acknowledge the support of UK’s Economic and Social Research Council/Medical Research Council grant (RES-060-25-0010), Biotechnology and Biological Sciences Research Council grant (BBE0039581), French Fondation pour la Recherche Médicale ( AJE201214), French Agence Nationale de la Recherche (ANR-16-CE37-0011-01 PRIMAVOICE; ANR-16-CONV-0002 Institute for Language, Communication and the Brain; ANR-11-LABX-0036 Brain and Language Research Institute), and the European Research Council (ERC) under the European Union’s Horizon 2020 research and innovation program (788240). The legacy STRAIGHT and its open sourcing was supported by the CREST program of the Japan Science and Technology Agency, the e-Society program of the Ministry of Education, Culture, Sports, Science, and Technology Japan, and the Advanced Telecommunications Research Institute International in Kyoto Japan.

**STRAIGHTMORPH source code and examples are available at: https://github.com/pascalbelin/STRAIGHTMORPH/**

**The authors declare no competing interests**

